# Intelligent Design of 14-3-3 Docking Proteins Utilizing Synthetic Evolution Artificial Intelligence (SYN-AI)

**DOI:** 10.1101/587204

**Authors:** Leroy K. Davis

**Affiliations:** Prairie View A & M University, Cooperative Agricultural Research Center (CARC), 700 University Drive, Prairie View, Texas 77446-0518

**Keywords:** Writing DNA code from scratch, Intelligent gene design, Synthetic evolution artificial intelligence, Time based DNA codes

## Abstract

The ability to write DNA code from scratch will allow for the discovery of new and interesting chemistries as well as allowing the rewiring of cell signal pathways. Herein, we have utilized synthetic evolution artificial intelligence (SYN-AI) to intelligently design a set of 14-3-3 docking genes. SYN-AI engineers synthetic genes utilizing a parental gene as an evolution template. Wherein, evolution is fast forwarded by transforming template gene sequences to DNA secondary and tertiary codes based upon gene hierarchical structural levels. The DNA secondary code allows identification of genomic building blocks across an orthologous sequence space comprising multiple genomes. Where, the DNA tertiary code allows engineering of super secondary structures. SYN-AI constructed a library of 10 million genes that was reduced to three structurally functional 14-3-3 docking genes by applying natural selection protocols. Synthetic protein identity was verified utilizing Clustal Omega sequence alignments and Phylogeny.fr phylogenetic analysis. Wherein, we were able to confirm three-dimensional structure utilizing I-TASSER and protein ligand interactions utilizing COACH and Cofactor. Conservation of allosteric communications were confirmed utilizing elastic and anisotropic network models. Whereby, we utilized elNemo and ANM2.1 to confirm conservation of the 14-3-3 ζ amphipathic groove. Notably, to the best of our knowledge, we report the first 14-3-3 docking genes to be written from scratch.

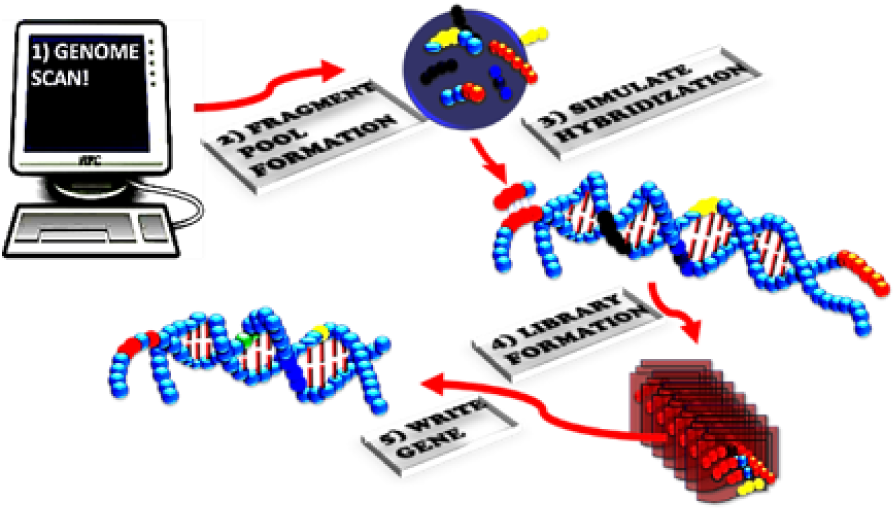

## 1. Introduction

The ability to write DNA code from scratch is a primary goal in the area of synthetic biology. Wherein, intelligent gene design will allow researchers to address a broad range of scientific conundrums such as the rewiring of signal pathways and the intelligent design of small genomes. Thusly, allowing for engineering of novel organisms and the potential for new and interesting chemistries that may include degradation of non-biodegradable products. Saliently, these technologies will open frontiers in medicine allowing the design of novel drug receptors for discovery of new cancer and disease treatments. The possibilities such a technology can offer are seemingly endless and essential for our current technological challenges. It is worth mentioning that over the past decade there has been multiple attempts at the modest ambition of de novo protein engineering. Wherein, the science was limited to a range of mutagenesis techniques that often resulted in non-functional proteins. While, there has been considerable progress the ability to intelligently design fully functional genes from scratch has been elusive.

In the current study, we focus on intelligent gene design utilizing synthetic evolution artificial intelligence (SYN-AI), an AI that accelerates the evolution process by performing a domain shuffling like mechanism similar to the ‘Domain Lego’ principle. ^1, 2, 3^ Evolutional acceleration is achieved by transforming gene sequences into DNA secondary (DSEC) and tertiary codes (DTER) based on gene hierarchical structure levels. We assume modern genes have a common ancestor that partitioned over time via DNA crossovers and that genetic diversity occurred by processes such as gene duplication, inversion, insertion and deletion. ^4, 5^ Our assumption of a common ancestor is in agreement with the “Universal Ancestor” and LUCA “Last Universal Common Ancestor” models. ^6, 7^ Saliently, the DSEC allows for identification of short highly conserved sequences occurring across multiple genomes referred to herein as genomic building blocks (GBBs). Based upon “*The Fundamental Theory of the Evolution Force*”, these sequences are genetic artifacts formed during the evolution process. ^71^ The DTER allows partitioning of genes at the super secondary structure level. Whereby, synthetic genes are engineered by walking the DTER followed by random selection and ligation of super secondary structures. Thusly, the exchange of information is analogous to the swapping of genomic building blocks in a game of Legos.

In order to fast forward the evolution process, SYN-AI forms an expanse orthologous sequence space followed by an exponential number of DNA crossovers within genomic alphabet comprising the DNA secondary code. SYN-AI identifies genomic building block formation across genomes by analysis of evolution force associated with DNA crossovers. Whereby, evolution force is a compulsion acting at the matter-energy interface that accomplishes genetic diversity while simultaneously conserving architecture and function. ^71^ In the current study, we utilize SYN-AI to intelligently design a set of 14-3-3 docking genes. These genes are responsible for regulating protein-protein interactions (PPI) in cell signal pathways. Saliently, docking protein interactions can have a profound effect on the target protein, altering its localization, stability, conformation, phosphorylation state, activity, and/or molecular interactions. ^9^ Thusly, the ability to write 14-3-3 docking genes from scratch will allow the rewiring of cell signal pathways. In addition to acting as key components of cell signal pathways 14-3-3 docking proteins play a role in cell growth and development, cancer cell signaling ^9, 10^, cellular metabolism and organelle communication. ^11^ Whereby, 14-3-3 proteins have been shown to interact with key signaling components such as the insulin like growth receptor ^57-59^, PI3K ^60, 61^, cdc25 phosphatase ^62-64^, and bad. ^65-67^.

In the current study, we intelligently designed a set of 14-3-3 docking genes utilizing SYN-AI. Wherein, a truncated *Bos taurus* 14-3-3 ζ docking gene was utilized as a template for gene engineering. Genomic building blocks were identified across multiple genomes by partitioning the parental docking gene into a DNA secondary code and simulating evolution by performing an exponential number of DNA crossovers within genomic alphabet forming the DSEC, **Fig.1** (top). Whereby, DNA crossover partners were randomly selected across an orthologous sequence space. Synthetic super secondary structures were engineered by targeting evolution within genomic alphabet comprising the DNA tertiary code, **Fig. 1** (middle). In all, SYN-AI generated a library of 10 million genes by walking the DTER, followed by random selection and ligation of synthetic super secondary structures, **Fig 1** (bottom). This expanse gene library was reduced to three genes utilizing natural selection protocols. Synthetic 14-3-3 docking genes were confirmed utilizing the Clustal Omega multiple sequence alignment tool and by phylogenetic analysis utilizing Phylogeny.fr. We were able to confirm three-dimensional structure of synthetic proteins utilizing I-TASSER as well as verify conservation of small molecule and fusicoccin binding sites utilizing Cofactor and Coach. Conservation of the 14-3-3 ζ amphipathic groove ^57^ as well as allosteric interactions were confirmed utilizing the elastic network model. Whereby, we utilized ElNemo and ANM2.1 to analyze normal modes, RMSD and deformation energies. Notably, based upon the aforementioned we confirm intelligent design of a set of novel 14-3-3 docking proteins utilizing synthetic evolution artificial intelligence.

**Figure 1.**
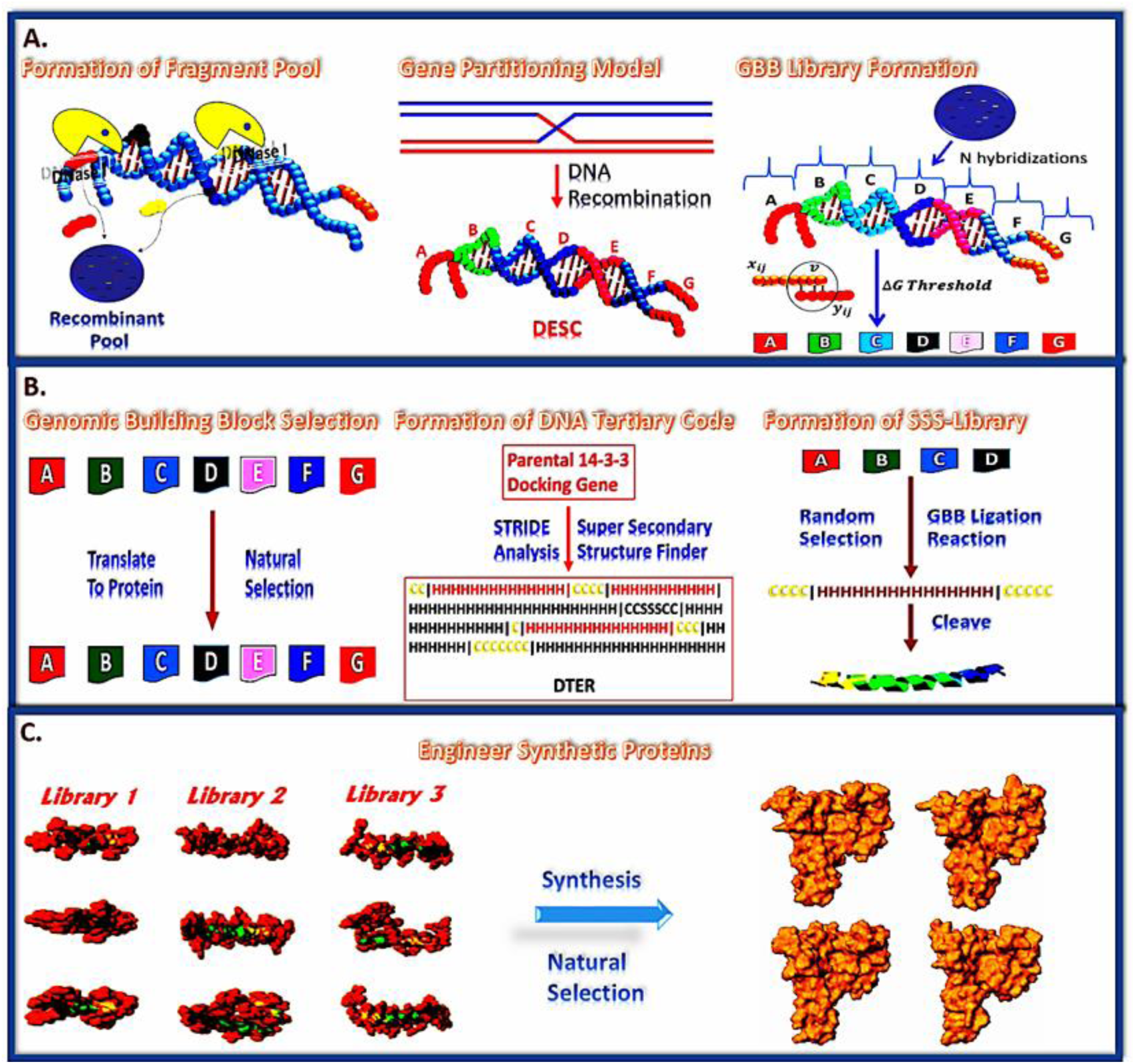
SYN-AI Mechanism. SYN-AI partitions the parental gene into a DNA secondary code (DSEC) and fast forwards the evolution process by performing an exponential number of DNA crossovers within each genomic alphabet comprising the DSEC, (**Top**). DNA crossovers characterized by the highest magnitude of evolution force are selected and stored in libraries. Subsequently, genomic building block (GBB) libraries are subjected to natural selection. Following natural selection, synthetic super secondary structures are formed by random selection and ligation of GBBs from appropriate genomic alphabet libraries, (**Middle**). Synthetic genes are engineered by walking the DNA tertiary code (DTER) followed by random selection and ligation of synthetic super secondary structures. Gene libraries are restricted to functional 14-3-3 docking genes by a natural selection process (**Bottom**).

## 2. Results

SYN-AI identified genomic building block formation by performing an exponential number of DNA crossovers within genomic alphabet forming the DSEC followed by analysis of the magnitude of evolution force associated with the aforementioned. Whereby, evolution force was analyzed over single and multidimensional planes of evolution formed as functions of the four evolution engines, i) evolution conservation, ii) wobble, iii) DNA binding state and (iv) periodicity according to “*The Fundamental Theory of the Evolution Force*” and as described in. ^71^ In addition, genomic building blocks were identified by applying natural selection protocols that limited selection to evolutionarily conserved DNA crossovers utilizing pattern recognition filters and that limited selection to sequences comprised of naturally occurring mutations utilizing Blosum 80 mutation frequency based algorithms. Subsequently, SYN-AI engineered synthetic 14-3-3 docking genes by walking the DNA tertiary code and randomly selecting and ligating synthetic super secondary structures formed by Domain Legos shuffling of genomic building blocks. Whereby, SYN-AI limited selection to structurally functional 14-3-3 docking genes by application of natural selection protocols.

Evolution force was analyzed utilizing the rotation model as described in ^71^ and as illustrated in **Fig. 2**. Wherein, genomic building blocks appeared as low density occurrences displaying (+) molecular wobble and high magnitude moments of inertia about the evolutional axis. In identifying genomic building blocks SYN-AI analyzed evolution force across single and multidimensional planes formed by the four evolution engines. Evolution force distribution across a one-dimensional evolution plane is illustrated in **Fig. 2**. Where, evolution force associated with genomic building block formation was analyzed within genomic alphabets (1-4) of the parental bovine 14-3-3 docking gene DNA secondary code.

**Figure 2.**
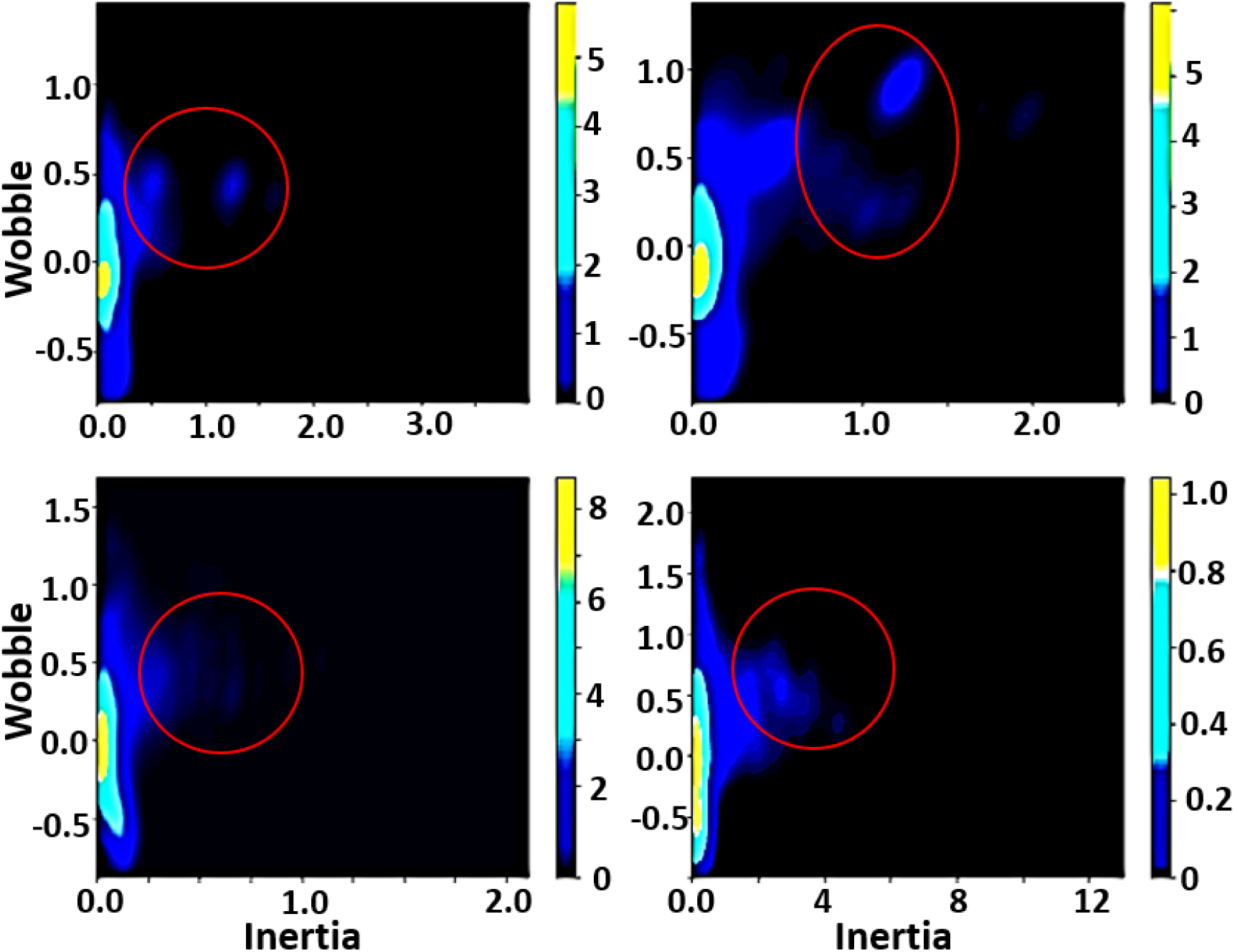
Analysis of Evolution Force Utilizing the Rotation Model. Evolution force was evaluated in genomic alphabets (1-4) of the parental bovine brain 14-3-3 docking gene DNA secondary code. Evolution force was evaluated as described in ^71^.

Protein identity was assessed by aligning synthetic docking genes with the truncated *Bos taurus* 14-3-3 ζ docking gene utilizing the Clustal Omega multiple sequence alignment tool. ^12^ Synthetic proteins displayed 58.5 to71.5 percent identity to the parental sequence characterized by stretches of high sequence identity between residues 100 – 130 and residues 154 – 180, **Fig. 3** (A). Phylogenetic analysis was performed utilizing Phylogeny.fr. ^13, 14^ Saliently, SYN-AI-1 and SYN-AI-2 diverged from the parental bovine gene forming novel 14-3-3 gene families. SYN-AI-3 was most closely related to the parental bovine 14-3-3 docking gene. However, each of the synthetic genes displayed significant branch distance from the parental gene, **Fig. 3** (B). We also performed phylogeny.fr blast, **Fig. 4**. Notably, each synthetic protein was characterized by distinctive phylogenetic relationships. Whereby, SYN-AI-1 was most closely related to 14-3-3 protein zeta/delta of *Ophiophagus hannah* and *Anolis carolinensis.* The aforementioned phylogenetic relationships were characterized by score: 281 bits (718), expect: 7e-73, identities: 150/213 (70%) and positives: 165/213 (77%). SYN-AI-2 also displayed close identity to the aforementioned but with greater divergence. Wherein, the relationship was characterized by alignment score: 266 bits (679), expect: 2e-68, identities: 142/212 (66%) and positives: 156/212 (73%). Saliently, SYN-AI-3 exhibited close phylogenetic relationship to protein zeta/delta of *Ovis aries* and the parental *Bos taurus* 14-3-3 ζ docking protein characterized by an alignment score: 281 bits (719), expect: 5e-73, identities: 149/213 (69%) and positives: 169/213 (79%).

**Figure 3.**
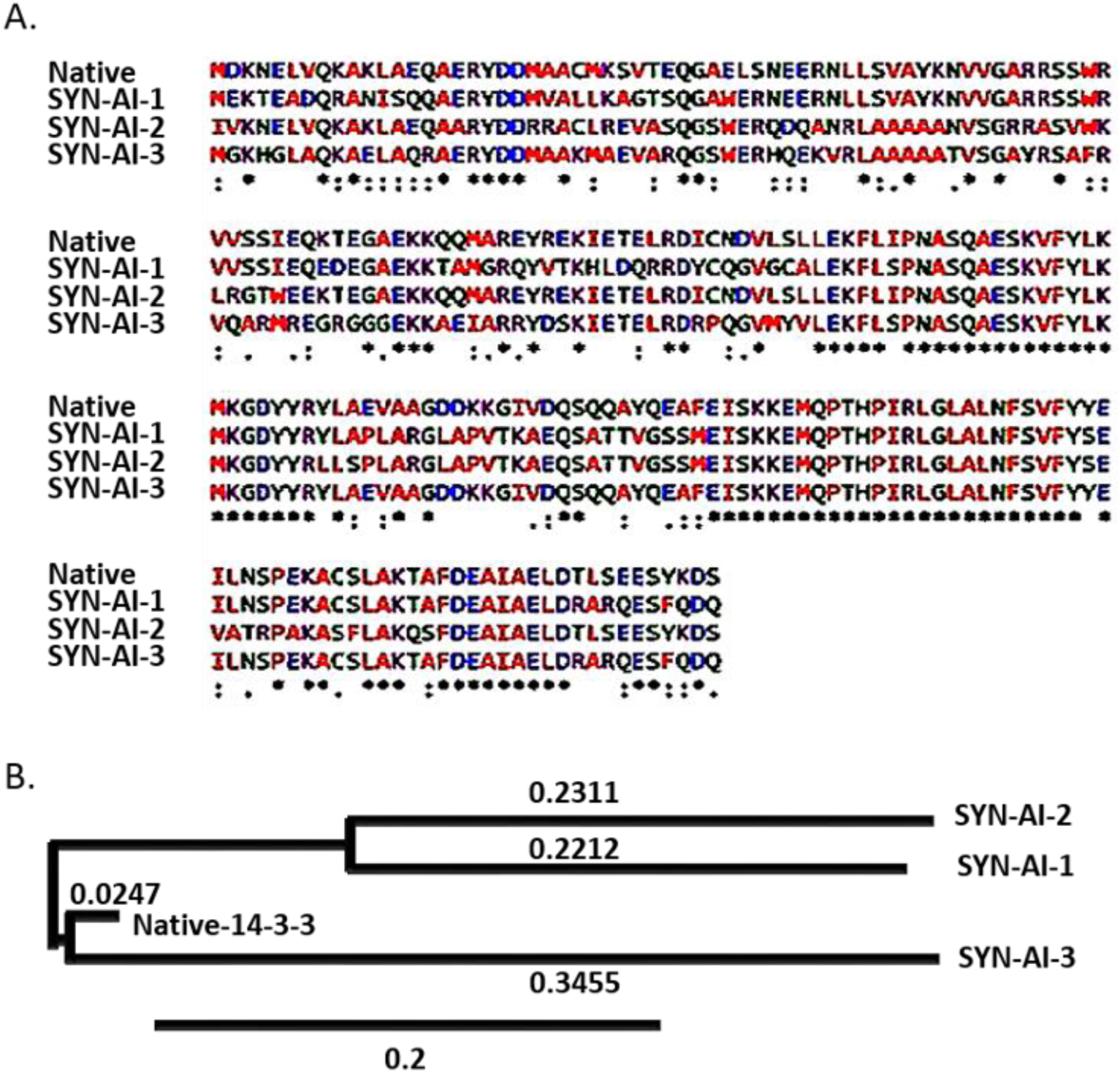
Sequence Alignment and Phylogenetic Analysis of Synthetic 14-3-3 Docking Proteins. Synthetic 14-3-3 docking proteins were aligned to truncated parental 14-3-3 ζ docking protein utilizing the Clustal Omega multiple sequence alignment tool, (**A**). Phylogenetic relationships between synthetic 14-3-3 docking proteins were compared to the parental utilizing Phylogeny.fr. (**B**).

**Figure 4.**
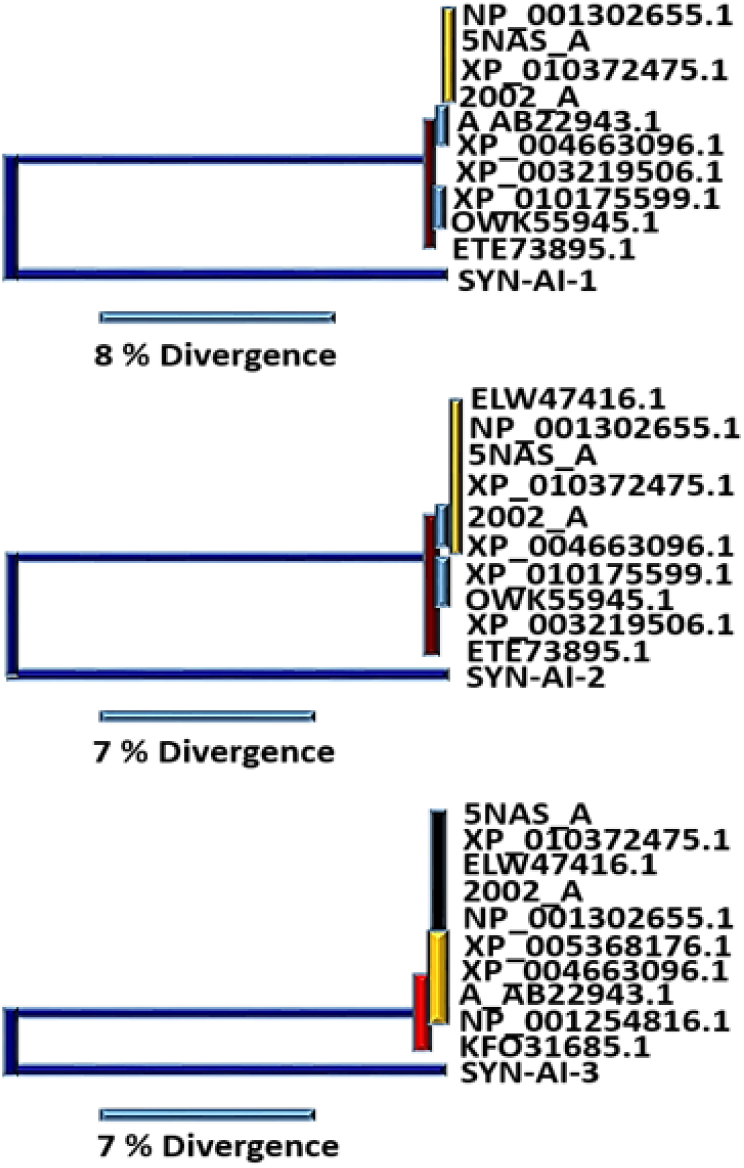
Phylogenetic Analysis of Synthetic Proteins. Phylogenetic relationships characterizing synthetic 14-3-3 docking genes engineered utilizing SYN-AI were analyzed by performing a Phylogeny.fr blast. Phylogenetic tree depicting synthetic protein SYN-AI-1 (**Top**), SYN-AI-2 (**Middle**) and SYN-AI-3 (**Bottom**).

Synthetic protein three-dimensional structure was analyzed utilizing the I-TASSER suite, Zhang Laboratory University of Michigan. Structural analysis revealed that synthetic proteins conserved 14-3-3 ζ architecture and surface as well as conserving volume of the ligand binding site, **Fig. 5**. SYN-AI-3 folded at highest confidence characterized by a C-score score of 1.54 and resolution of 2.5 ± 1.9Å. However, I-TASSER structural predictions of all three synthetic proteins were very reliable wherein synthetic proteins folded at an average C-score of 1.51. Saliently, the truncated parental 14-3-3 ζ docking protein folded at a similar confidence score of 1.55 with a TM-score of 0.93±0.06 and a RMSD of 2.5 ± 1.9Å. In addition, SYN-AI-1 and SYN-AI-3 were characterized by a TM-score of 0.93±0.06 compared to SYN-AI-2 that was characterized by a TM-Score of 0.92±0.06. Further, SYN-AI-1 and SYN-AI-3 were predicted at an RMSD of 2.5±1.9Å wherein SYN-AI-2 was predicted at an RMSD of 2.6±1.9Å.

**Figure 5.**
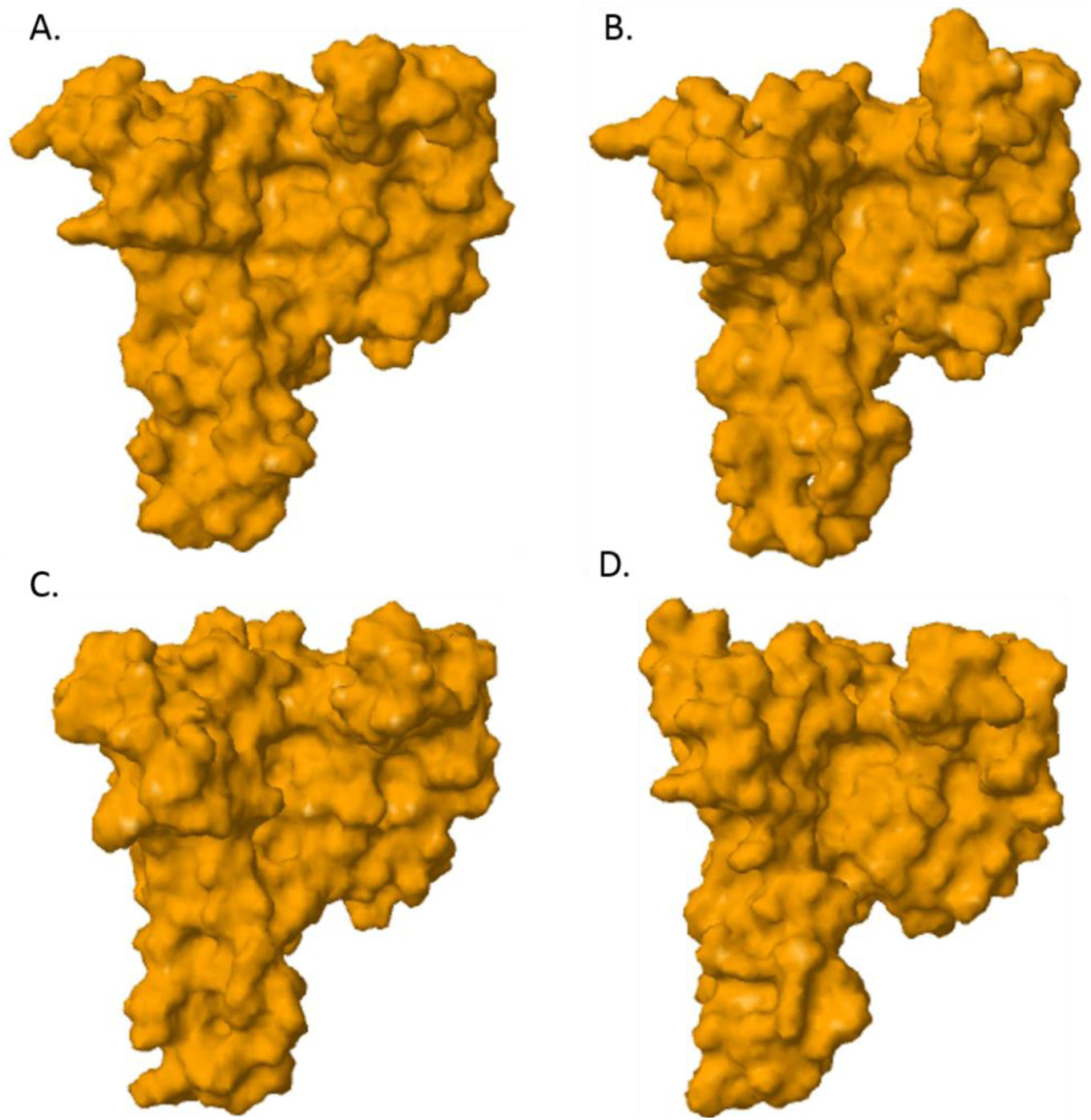
Three-dimensional Structure Predictions. The I-TASSER Suite, Zhang Laboratory University of Michigan was utilized to analyze three-dimensional structure. Molecular surface image of truncated parental 14-3-3 ζ docking protein predicted at a confidence of 1.55 and a resolution of 2.5 ± 1.9 Å, **(A**). Molecular surface image of SYN-AI-3 predicted at a confidence of 1.54 and a resolution of 2.5 ± 1.9 Å, (**B**). Molecular surface image of SYN-AI-1 predicted at a confidence of 1.51 and a resolution of 2.5 ±1.9 Å, (**C**). SYN-AI-2 predicted at a confidence of 1.48 and a resolution 2.6 ± 1.9 Å, (**D**).

Synthetic protein ligand binding interactions were analyzed utilizing Cofactor and Coach. Notably, SYN-AI successfully conserved fungal toxin fusicoccin complex (FC) binding within the BS03 site of SYN-AI-3, **Fig. 6** (top Left). The aforementioned was predicted at a confidence score of 0.40 and a resolution of 2.74 Å. Wherein, FC binding within parental bovine 14-3-3 docking protein was predicted at a slightly higher confidence of 0.45. The aforementioned analysis revealed that fusicoccin ligand-residue interactions were conserved with exception of N42 → V42 and V46 → A46 mutations. Fusicoccin formed hydrogen bonding interactions with residues V42, A46, K120, M121, P165, I166 and D213 and Van der Waals interaction with F117 of the synthetic protein amphipathic groove **Fig. 6** (Top Right). Whereby, the FC complex formed bonding interactions with residue V42 at a distance of 0.285 nm, A46 at 0.375 nm, F117 at 0.223 nm, K120 at 0.357 nm, M121 at 0.306 nm, P165 at 0.332 nm, I166 at 0.300 nm and residue D213 at 0.353 nm. Compared to parental 14-3-3 docking protein, where fusicoccin formed bonding interactions with residue N42 at a distance of 0.41 nm, V46 at 0.353 nm, F117 at 0.267 nm, K120 at 0.322 nm, M121 at 0.333 nm, P165 at 0.332 nm, I166 at 0.285 nm and residue D213 at a distance of 0.42 nm. Notably, the FC complex ligand binding site was fully conserved in SYN-AI-1, **Fig. 7**. Where, fusicoccin formed bonding interactions with residue N42 at a distance of 0.354 nm, V46 at 0.27 nm, F117 at 0.213 nm, K120 at 0.333 nm, M121 at 0.287 nm, P165 at 0.33 nm, I166 at 0.379 nm and D213 at a distance of 0.386 nm. Comparatively, synthetic protein SYN-AI-2 conserved the fusicoccin ligand binding site with the exception of a V46 → A46 mutation, **Fig. 7**. Whereby, fusicoccin formed bonding interactions with residue N42 at a distance of 0.222 nm, A46 at 0.445 nm, F117 at 0.282 nm, K120 at 0.320 nm, M121 at 0.344 nm, P165 at 0.329 nm, I166 at 0.301 nm, and D213 at a distance of 0.343 nm.

**Figure 6.**
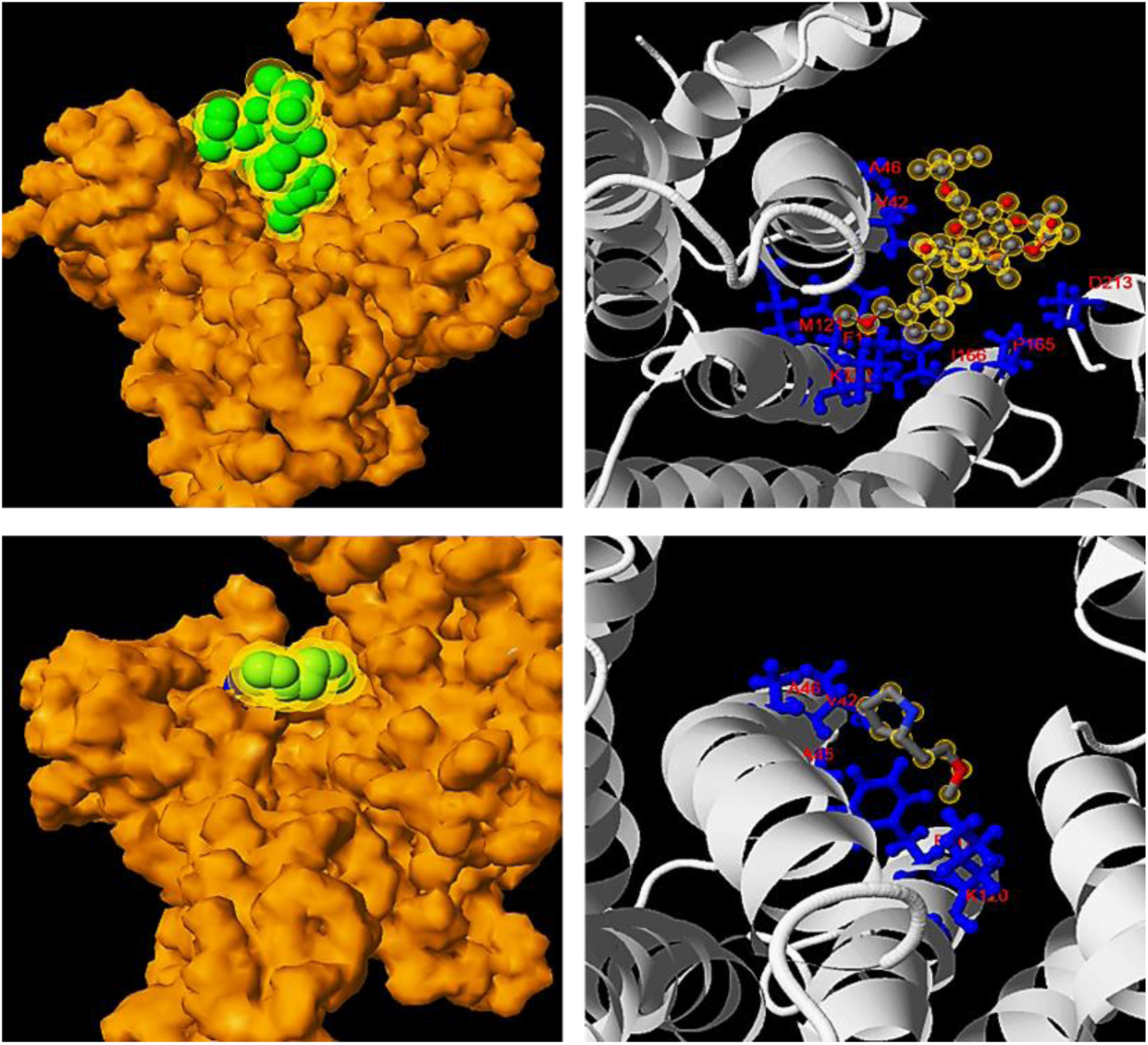
Analysis of Protein-ligand Interactions. Ligand and small molecule interactions were analyzed utilizing Cofactor and Coach. Van der Waals surface image of fusicoccin binding at a resolution of 2.74 Å, (**Top Left**). Ligand residue interactions between fusicoccin and SYN-AI-3 (**Top Right**). Van der Waals surface image of small molecule (2S)-(2)-methoxyethyl pyrrolidine at a resolution of 1.7 Å, (**Bottom Left**). Ligand residue interactions between SYN-AI-3 and (2S)-(2)-methoxyethyl pyrrolidine (**Bottom Right**).

**Figure 7.**
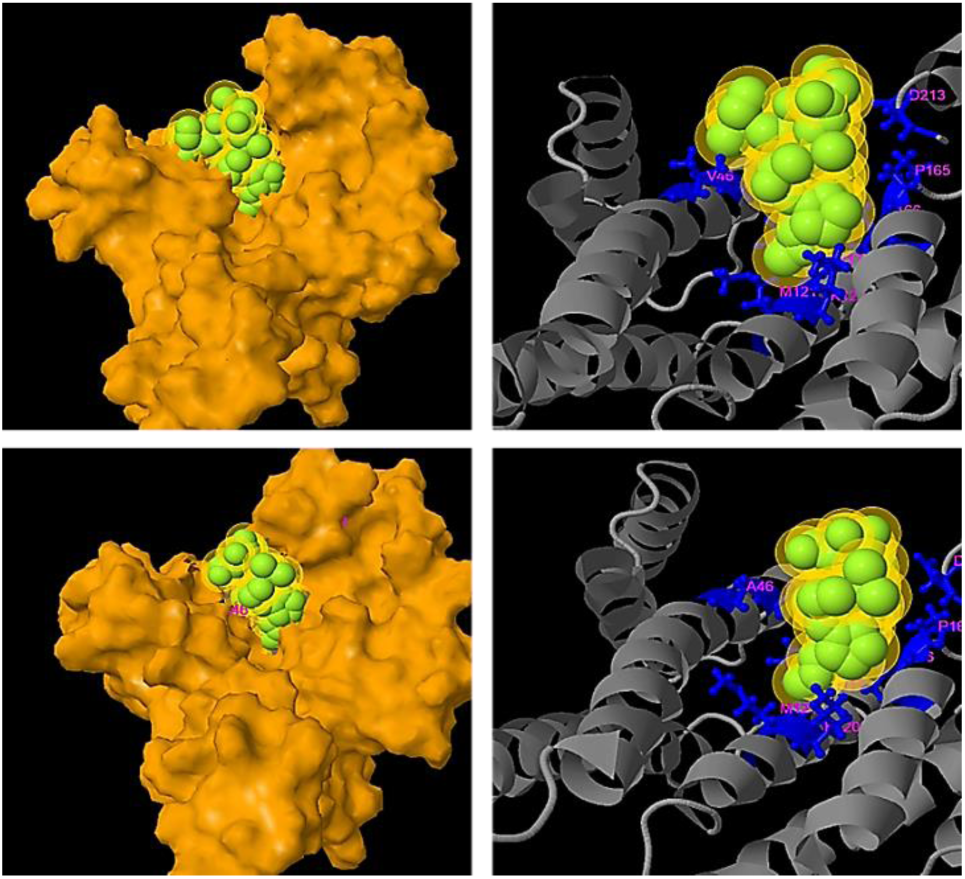
Analysis of Ligand Binding Interactions. Synthetic protein ligand binding interactions were analyzed utilizing Cofactor and Coach. Molecular surface image of fusicoccin SYN-AI-1 ligand binding reaction predicted at a C-score of 0.46, (**Top Left**). SYN-AI-1 fusicoccin ligand-residue interactions (**Top Right**). Molecular surface image of fusicoccin SYN-AI-2 ligand binding interaction predicted at a C-score of 0.41, (**Bottom Left**). SYN-AI-2 fusicoccin ligand-residue interactions, (**Bottom Right**).

In addition to conserving FC complex ligand residue interactions, synthetic evolution artificial intelligence conserved small molecule (2S)-2-(2-methoxyethyl) pyrrolidine ligand binding within the BS02 site, **Fig. 6** (Bottom). Wherein, Cofactor and Coach predicted binding of the small molecule within the SYN-AI-3 BS02 site at a resolution of 1.7 Å. Our analysis revealed that the SYN-AI-3 BS02 ligand binding site comprised N42 → V42, S45 → A45 and V46 → A46 mutations. Further, analysis of ligand-residue interactions within the SYN-AI-3 BS02 site revealed that (2S)-2-(2-methoxyethyl) pyrrolidine interacts with residue V42 at a distance of 0.264 nm, A45 at 327 nm, A46 at 0.391 nm, F117 at 0.252 nm and residue K120 at a distance of 0.397 nm. Comparatively, (2S)-2-(2-methoxyethyl) binding within the parental bovine 14-3-3 docking protein BS02 site was characterized by interaction of residues N42 at 0.244 nm, S45 at 0.305 nm, V46 at 0.268 nm, F117 at 0.282 nm, and residue K120 at a distance of 0.366 nm. Notably, synthetic protein SYN-AI-1 displayed full conservation of the BS02 ligand binding site. Whereby, (2S)-2-(2-methoxyethyl) pyrrolidine interacted with residue N42 at a distance of 0.261 nm, S45 at 0.273 nm, V46 at 0.270 nm, F117 at 0.250 nm, and residue K120 at a distance of 0.385 nm.

Notably, synthetic evolution artificial intelligence successfully conserved protein-protein interactions as corroborated by Coach and Cofactor analysis of BS01 and BS02 ligand binding sites. Whereby, test probe I (TMLNLVSGRRR) occupied and was deeply buried within BS01 and BS02 ligand binding sites, **Fig. 8** (Top Left). Test probe I ligand interaction was predicted at 2.74 Å with a C-score of 0.31. Whereby, the probe interacted with residues H38, K41, V42, A45, A46, R56, R60, F117, K120, R127, Y128, P165, I166, G169, L172, N173, V176, and E180 of the synthetic protein amphipathic grove **Fig. 8** (Top right). Test probe II (VTYSG) binding within the BS01 site was predicted at a C-score of 0.82 and at a resolution of 2.3 Å, **Fig. 8** (Bottom Left). Whereby, probe II interacted with residues K49, R56, R60, K120, R127, Y128, L172, V176 and E180 of the amphipathic grove, **Fig. 8** (Bottom Right). Saliently, synthetic evolution artificial intelligence accomplished full conservation of BS01 ligand-binding residues within synthetic protein SYN-AI-3 with the exception of a K49 → A49 mutation as well as an additional G169 residue contact. Comparatively, synthetic protein SYN-AI-1 displayed full conservation of ligand-residue interactions with addition of the G169 residue contact. Whereby, SYN-AI-2 comprised K49 → A49 and R56 → A56 mutations in addition to the G169 contact.

**Figure 8.**
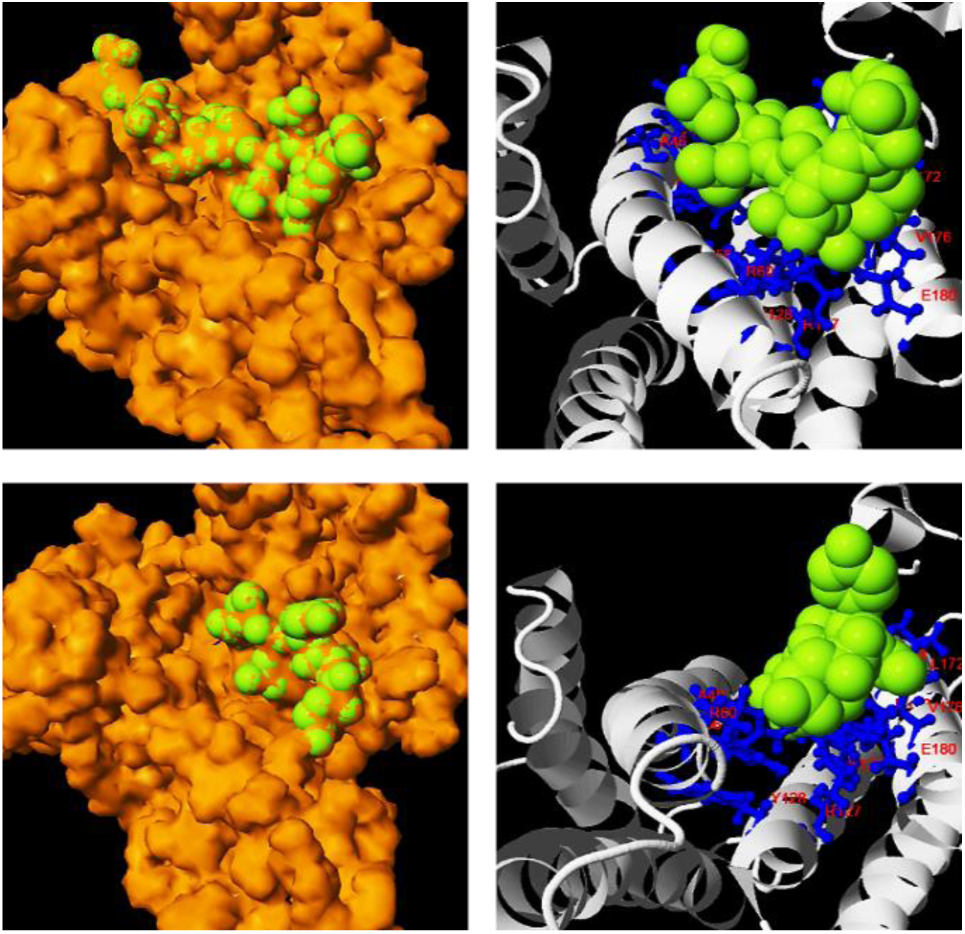
Analysis of Protein-protein Interaction Sites. Protein interactions were analyzed utilizing Cofactor and Coach. Van der Waals surface image of probe I (TMLNLVSGRRR) buried within BS01 and BS02 ligand binding sites, predicted at 2.74 Å (**Top Left**). Probe I ligand-residue interactions (**Top Right**). Van der Waals surface image of probe II (VTYSG) buried within the BS01 site, predicted at 2.3 Å, (**Bottom Left**). Probe II ligand-residue interactions, (**Bottom Right**).

In order to demonstrate SYN-AI’s ability to conserve protein allosteric effects, we analyzed the parental and SYN-AI engineered 14-3-3 ζ docking proteins utilizing the elastic network model. Whereby, truncated parental and synthetic structures were predicted utilizing I-TASSER. ElNemo ^68, 69^ was utilized to perform normal mode analysis, **Fig. 9**. Normal mode analysis resulted in 107 modes of which five (7-11) were low frequency modes indicating a role in ligand binding and intra protein communication. Saliently, Cα strains within synthetic proteins were similar to those occurring within the parental docking protein as shown in **Fig. 9** (A). Whereby, mode7 was characterized by a mean residue sample variance of *σ*^2^ = 2.62 × 10^-4^. Residue root mean square deviations of synthetic proteins also closely overlapped the parental 14-3-3 ζ docking protein, **Fig. 9** (B). When comparing RMSD of parental 14-3-3 ζ docking protein to synthetic proteins there exists a miniscule variance of *σ*^2^ = 1.115 × 10^−4^ Å. Normal mode analysis also revealed that the frequency and collectivity of synthetic protein modes closely mirrored those of the parental 14-3-3 ζ docking protein, **Fig. 9** (C, D). Frequency of synthetic modes were characterized by a variance of *σ*^2^ = 0.202 from the parental docking protein. Wherein, synthetic proteins were characterized by a variance in collectivity of *σ*^2^ = 9.593 × 10^−3^ from parental 14-3-3 ζ. The anisotropic network model ANM2.1^70^ was utilized to analyze energy deformation and solvent accessibility. Synthetic protein energy deformation peak pattern and strength were analogous to that of the parental protein with a strong deformation peak ranging from residues 43-106 as well as residue 127-148, **Fig. 10** (A, B, C, D). Notably, energy deformation peaks correlate well with the location of the 14-3-3 ζ amphipathic groove as well as ligand binding residues predicted by I-TASSER. Wherein, the strong peak at residue 127 suggests a significant role of the Van der Waal interaction with arginine in ligand binding, in both the parental and synthetic proteins. Synthetic protein solvent accessibility also overlapped with that of the parental protein with little variation per residue as illustrated in **Fig. 10** (E).

**Figure 9.**
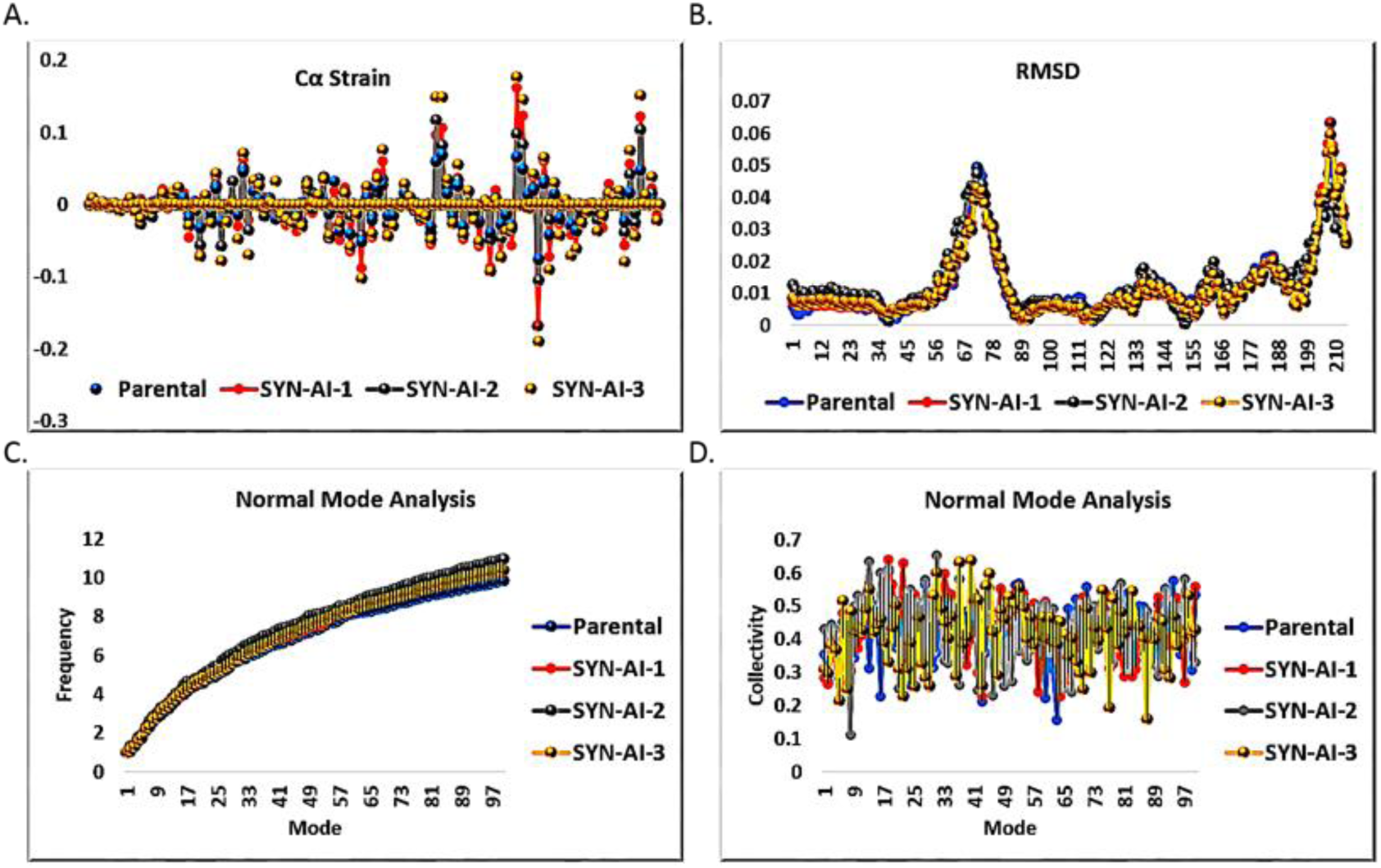
Analysis of Allosteric Interactions. ElNemo was utilized to perform normal mode analysis of parental 14-3-3 ζ and synthetic proteins SYN-AI-1, SYN-AI-2 and SYN-AI-3. Whereby, we analyzed carbon alpha strain (**A**), root mean square deviation (**B**), mode frequency (**C**), mode collectivity (**D**).

**Figure 10.**
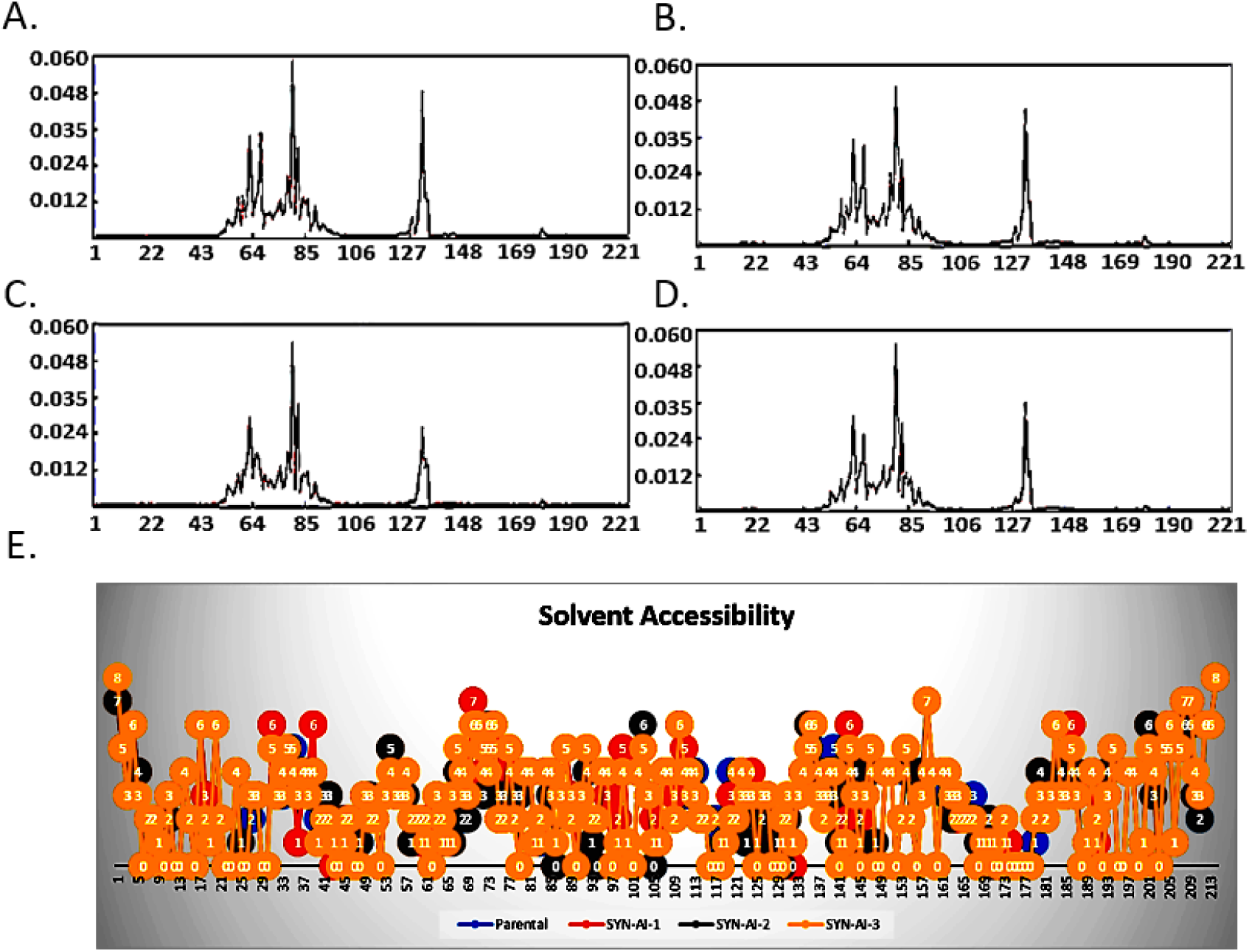
Analysis of Amphipathic Groove. Conservation of the 14-3-3 ζ amphipathic groove was confirmed by the anisotropic model as described in ^70^. Whereby, ANM2.1 was utilized to analyze energy deformations occurring in the parental 14-3-3 ζ docking protein (**A**), SYN-AI-1(**B**), SYN-AI-2 (**C**) and synthetic protein SYN-AI-3 (**D**). To further verify conservation of the amphipathic groove, ANM2.1 was also utilized to predict parental and synthetic 14-3-3 ζ docking protein solvent accessibility (**E**).

## 3. Discussion

In the current study, we utilized SYN-AI to design a set of 14-3-3 docking proteins utilizing *Bos taurus* 14-3-3 ζ docking protein as a parental template. Notably, SYN-AI is not a rational design technology, but simulates evolution by evaluating evolution force associated with genomic building block formation and subsequently builds genes from scratch by randomly assembling the aforementioned. Our approach anticipates the evolution process by simulating DNA shuffling. Thusly, SYN-AI is advantageous in engineering functional genes as anticipatory evolution has been shown to generate functional proteins with high efficacy ^74^. Evolution force was solved by overlapping gene sequences occurring over an orthologue sequence space with that of the parental template. Thereby, SYN-AI is able to analyze evolutional character of DNA crossovers going back to “LUCA”. It is worth mentioning that our technology is limited by its dependence on the availability of PDB structural data. In the current study, crystal data was available for 228 of 245 *Bos taurus* 14-3-3 ζ residues. Further, these crystal data contained gaps wherein STRIDE structure based analysis was able to generate structural data for 213 of the 228 residues. Thusly, SYN-AI protein engineering was limited to the available empirical data, whereby we showed the potential of our technology for gene and signal pathway engineering by synthesizing a set of truncated 14-3-3 docking proteins of 214 residues.

According to “*The Fundamental theory of the Evolution Force*” we were able to analyze evolution force associated with genomic building block formation based upon four evolution engines, (i) evolution conservation, (ii) wobble, (iii) DNA binding state and (iv) periodicity. Molecular biologists have long utilized evolution conservation as a tool when selecting mutable gene regions. Where, it has been assumed that highly evolutionarily conserved residues are critical to protein function. ^15-23^ Thusly, SYN-AI is based upon the hypothesis that evolution conservation is an artifact of the evolution force. Classically, wobble has been defined by genetic diversity in the third codon with conservation of residue sequence. ^24-30^ However, in fingerprinting the evolution force we expanded the definition of wobble to the acquisition of genetic diversity with conservation of architecture. Thusly, allowing wobble to be viewed at all structural levels. For instance, super secondary structures such as helices, turns and beta sheets are expressed in genetically diverse species yet retain basic architecture and function. A further example of wobble at a higher structural level is that of bipedal animals. Although the aforementioned are genetically diverse anatomical structure and architecture are reserved across species. Also when constructing SYN-AI, we assumed that evolution force interacts at the matter-energy interface via DNA crossovers. Thusly, we were able to evaluate interaction of the evolution force at the matter-energy interface by analyzing DNA binding states. Wherein, DNA binding states measure DNA crossover selectivity in respect to the recombinant pool and are a function of Gibb’s free energy associated with DNA base stacking interactions. ^8, 31-33^ The final evolution engine we considered was periodicity. We assumed that nature has a tendency to repeat successful structures that promote the survival of an organism. Thereby, evolution force at the molecular level is a function of sequence periodicity. ^34-36^

In the current study, we have demonstrated that genomic building blocks can be identified across multiple genomes by analyzing evolution force and exploited to write genes from scratch. Wherein, by transforming a parental gene template into DNA secondary and tertiary codes based on hierarchical structure levels, SYN-AI was able to fast forward the evolution process. The aforementioned allowed identification of genomic building blocks by characterization of evolution force and allowed for intelligent gene design by a Legos like swapping of genetic material in agreement with the ‘Domain Lego’ principle. ^1, 2, 3^ Notably, SYN-AI generated synthetic proteins displaying high sequence identity to naturally occurring 14-3-3 docking proteins suggesting that the AI successfully simulated the evolution process allowing divergence from the parental 14-3-3 ζ docking gene while evolutionarily conserving 14-3-3 global architecture. We were able to corroborate the aforementioned by performing phylogeny.fr blast and established that each synthetic protein comprised of diverse phylogenetic relationships as characterized by diverse phylogenetic trees. Synthetic proteins also displayed significant branch distance from one another further confirming that SYN-AI fast forwarded the evolution process. Wherein, each synthetic protein diverged into a different evolutional pathway. Saliently, despite performing ∼300 million DNA crossovers within genomic alphabet comprising the 14-3-3 DNA secondary code, Clustal Omega sequence alignments of synthetic proteins were characterized by stretches of high sequence identity wherein no genetic diversity was accomplished by SYN-AI. Notably, the aforementioned corroborates that our natural selection protocols implemented into SYN-AI successfully conserved slow evolving regions of genes. Saliently, resistance of these regions to mutation suggests they are essential to cellular function. ^37-39^

In addition to fast forwarding the evolution process, SYN-AI successfully conserved 14-3-3 docking protein architecture. Whereby, I-TASSER three dimensional structural analysis revealed that global 14-3-3 docking protein architecture was conserved in all three synthetic proteins. Based upon molecular dynamics simulation data in conjunction with previously discussed phylogenetic data, we validate our hypothesis of evolution force effects on genomic building block formation by proof of concept. Further corroborating our proof of concept, SYN-AI evolutionarily conserved ligand binding sites and ligand-residue interactions. Notably, analysis by Coach and Cofactor ^40^ confirmed that SYN-AI conserved small molecule binding as demonstrated by conservation of (2S)-2-(2-methoxyethyl) pyrrolidine binding within the BS02 site. While conserving small molecule binding synthetic evolution accomplished significant modification of the SYN-AI-3 BS02 ligand binding site characterized by mutation of three of the five residues participating in the binding of (2S)-2-(2-methoxyethyl) pyrrolidine. Thusly, SYN-AI successfully altered positioning and conformation of the molecule within the binding site. Contrary to SYN-AI-3, synthetic protein SYN-AI-1 was characterized by full conservation of the BS02 binding site. Wherein, the artificial intelligence preserved parental ligand-residue interactions. Saliently, conformation and positioning of (2S)-2-(2-methoxyethyl) pyrrolidine within the SYN-AI-1 BS02 ligand binding site was altered due to changes in volume resulting from the modification of global protein architecture due to the evolution process. The aforementioned is corroborated by phylogenetic analysis. Wherein, SYN-AI-1 is significantly diverged from parental bovine brain 14-3-3 docking protein in comparison to synthetic protein SYN-AI-3. Notably, the ability of synthetic evolution artificial intelligence to evolutionarily conserve small molecule binding sites while altering conformation and binding affinities of ligands is significant as small molecules stabilize and inhibit 14-3-3 protein-protein interactions that play a role in neurodegenerative diseases and cancer. ^41^

Contrary to the surprising level of divergence achieved in the BS02 ligand binding site, SYN-AI evolutionarily conserved amphipathic interfaces and ligand binding residues within the BS01 and BS03 ligand binding sites. Whereby, Coach and cofactor confirmed conservation of fusicoccin binding within all three synthetic proteins engineered utilizing synthetic evolution artificial intelligence. Fusicoccin ligand-residue interactions were also conserved with the exception of N42 → V42 and V46 → A46 mutations. In agreement with our previous experiments, modifications in global protein architecture altered conformation of fusicoccin within the amphipathic groove signifying that fusicoccin exhibited altered binding affinity for the ligand binding site. Notably, SYN-AI’s ability to evolutionarily conserve the FC complex binding site is significant in that the complex is responsible for activating *H*^+^ pumping across the plasma membrane. ^42^ In plants FC complex interactions with the 14-3-3 docking protein activates KAT1 channels and is responsible for cell growth by regulating diffusion through *K*^+^ channels.^43^ Saliently, the FC 14-3-3 complex also regulates defense responses in Tomato plants.^44^ In addition to conservation of the BS02 and BS03 sites, analysis of protein probe localization confirmed conservation of the BS01 ligand binding site with exception of a K49 → A49 mutation and an additional G169 residue contact. The aforementioned corroborates evolutional conservation of protein-protein interactions in synthetic proteins.

Notably, altered positioning and conformation of small molecules, protein probes and the FC complex within synthetic protein BS01, BS02 and BS03 sites substantiate our assumption that synthetic proteins designed in this study display the potential for altered PPI. The aforementioned is significant in that interaction of globular domains of protein interaction partners within the 14-3-3 amphipathic grove regulate stress signaling proteins such as ERK, MAPK, JNK and p38 MAPK as well as growth and cell cycle regulators raf, PI3K and cdc25 phosphatase. ^45-49, 61-67^ We confirmed conservation of these ligand binding interactions as well as confirmed the conservation of intra protein communication by analyzing allosteric effects utilizing the elastic network model. Whereby, normal mode analysis of parental and synthetic 14-3-3 ζ docking proteins was performed utilizing ElNemo and ANM2.1. Analysis of mode 7 indicated that there existed little variance in Cα strain occurring in parental 14-3-3 ζ and synthetic proteins. These data suggests that synthetic proteins are energetically stable and corroborate the validity of I-TASSER structure prediction. The aforementioned are corroborated by RMSD results, wherein synthetic proteins exhibited little variance from the parental 14-3-3 ζ protein. Thusly, SYN-AI was able to evolve protein sequence and local structures without disrupting global protein architecture. Notably, the aforementioned was accomplished while conserving mode frequency and collectivity suggesting that there exist little variance in residue potential energies during structural transitioning from parental to synthetic 14-3-3 ζ proteins. This is prominent due to the many ligand and signal pathway interactions occurring within the 14-3-3 ζ amphipathic groove as corroborated by the presence of 107 modes indicated by ElNemo. Our data suggests that SYN-AI conserved allosteric interactions regulating signal transduction pathways. Conservation of the 14-3-3 ζ amphipathic groove is corroborated by solvent accessibility data indicating only infinitesimal differences in protein hydrophobicity. Notably, we demonstrate conservation of the binding of the R18 protein peptide within the amphipathic groove as well as conservation of a “hug and squeeze” mechanism. Where, the left and right torso of the 14-3-3 monomer flex closed and secure the R18 protein peptide in the amphipathic groove. The open configuration is shown is **Fig. 11**(A, C) and the closed configuration in **Fig. 11** (B, D). Notably, we also successfully dimerized the SYN-AI-1 monomer and engineered a functional 14-3-3 ζ dimer, while maintaining the “bend and flex” mechanism present in the WT dimer as illustrated in **Fig. 12**. Whereby, we show the synthetic 14-3-3 ζ dimmer in both the open and closed positions. Modes 7-11 were also conserved in SYN-AI generated dimmers. We have demonstrated that SYN-AI conserved allosteric communications in the monomeric form of the 14-3-3 ζ protein as well as low frequency vibrations occurring in the 14-3-3 ζ dimeric form, thusly validating that identification of genomic building blocks by characterization of evolution force and synthetic gene evolution utilizing SYN-AI is an excellent tool for signal transduction pathway engineering.

**Figure 11.**
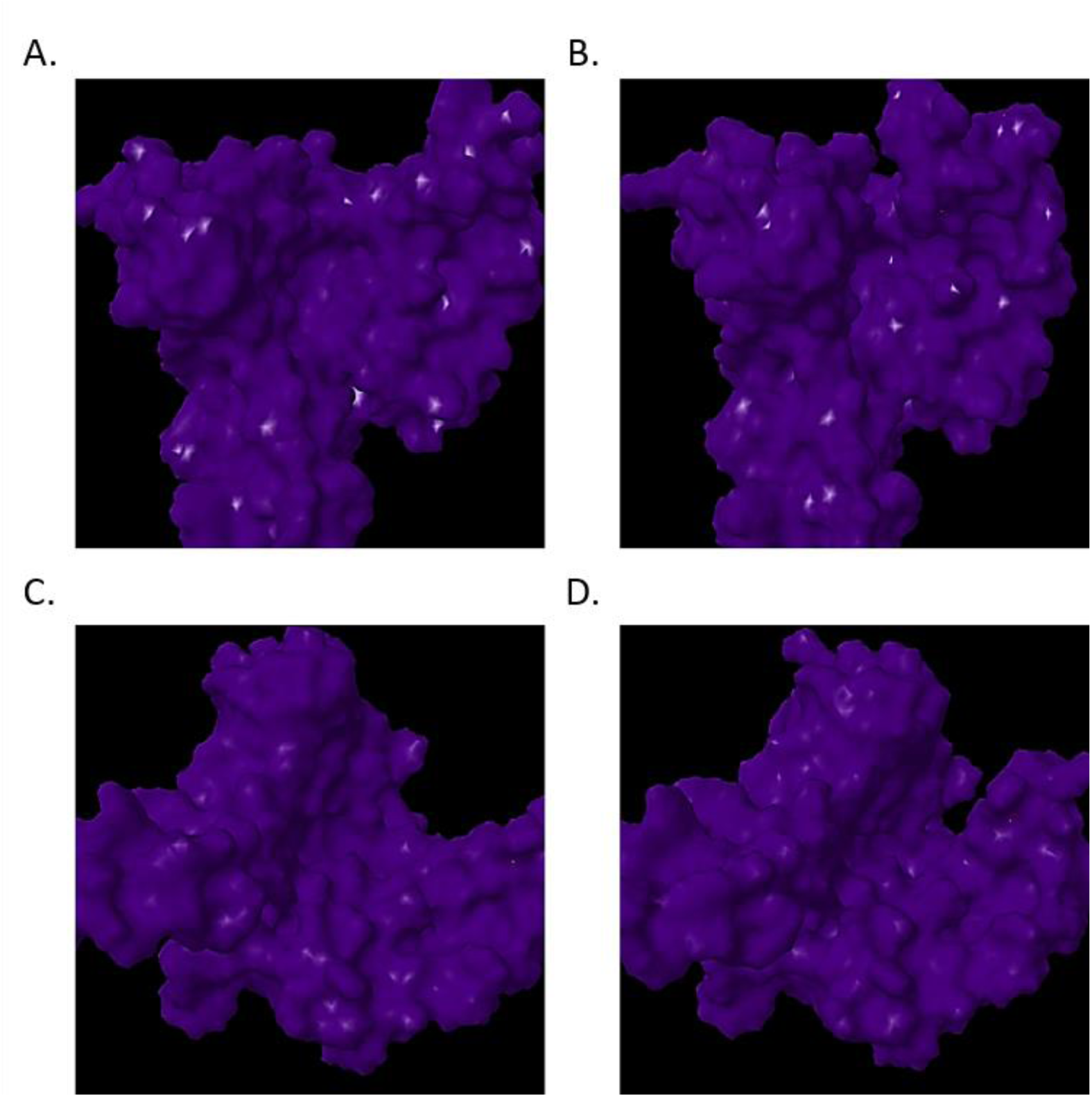
Hug and Squeeze Mechanism. The elastic network model was utilized to perform normal mode analysis. Low frequency vibrations occurring within parental and synthetic 14-3-3 ζ docking proteins were analyzed utilizing ElNemo. Frontal view of mode 7 open configuration (**A**), frontal view of mode 7 closed configuration (**B**), view of open configuration down amphipathic groove (**C**), view of closed configuration down amphipathic groove (**D**).

**Figure 12.**
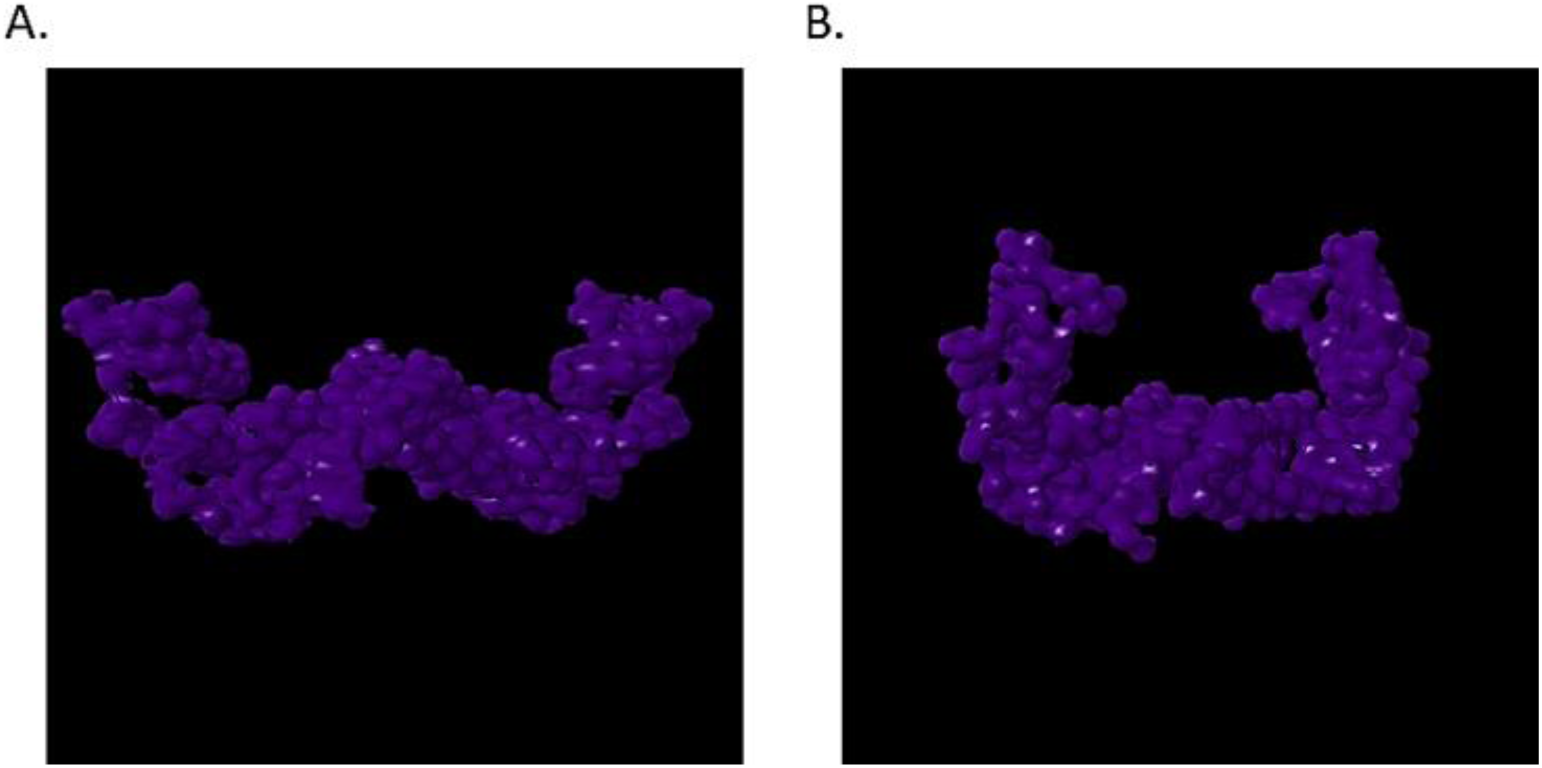
Synthetic 14-3-3 ζ Dimer Formation. Dimerization of the SYN-AI-1 14-3-3 monomer was predicted utilizing COTH ^73^, Zhang Laboratory University of Michigan. Normal mode analysis was performed utilizing ElNemo and graphics generated utilizing the Jena3D Viewer. The open configuration of the Mode 7 “bend and flex” mechanism is illustrated in (**A**), whereby the closed configuration is shown in (**B**).

Saliently, our study relies heavily on the accuracy of computational structure prediction. However, our results are corroborated by the parental 14-3-3 ζ structure reported in ^57^. Whereby, as an experimental control we overlapped the native 14-3-3 ζ crystal structure with the I-TASSER predicted PDB structure. As illustrated in **Fig. 13**, the I-TASSER 14-3-3 ζ structure prediction is identical to the reported crystal structure. Wherein, we utilized the SuperPose server ^72^ to overlap the PDB structure reported in ^57^ to the I-TASSER predicted structure, with only minor deviations present in the coiled coil regions. It is worth mentioning, this level of accuracy is not guaranteed for all subsequent predictions performed in this study. However, the claims presented herein are supported by sufficient scientific data to corroborate the experimental controls and I-TASSER structure prediction of synthetic proteins. Wherein, we can confidently conclude that SYN-AI successfully engineered a set of functional 14-3-3 ζ docking genes from scratch.

**Figure 13.**
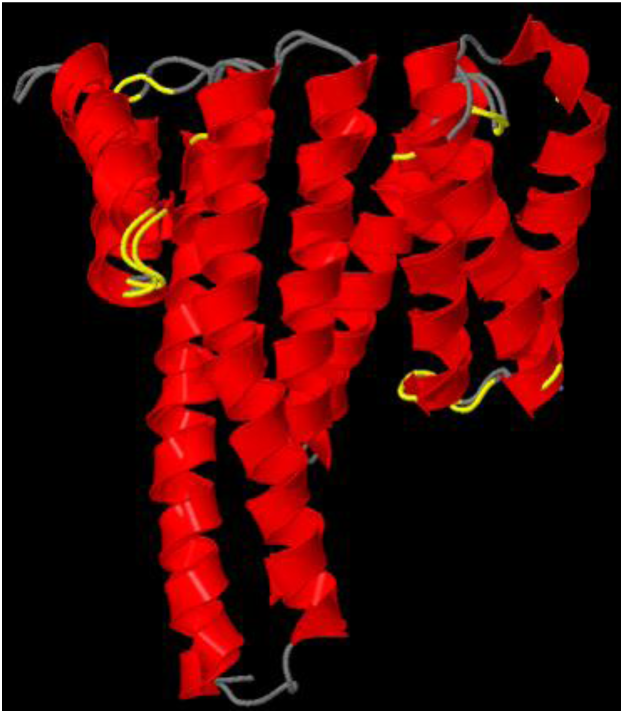
Superimposition of experimental and predicted 14-3-3 ζ structures. The SuperPose server was utilized to compare the native *Bos taurus* 14-3-3 crystal structure reported in ^57^ to the I-TASSER predicted structure. The native crystal structure covered residues 1-228 with two gaps of 5 and 7 amino acids, respectively. Superimposition of the parental and synthetic proteins was performed with a minimum sequence similarity of 25% identity, a similarity cutoff of RMSD of 2.0 Å, a dissimilarity cutoff of RMSD of 3.0 Å, and a dissimilar subdomain cutoff of 7 residues.

## Conclusion

Based upon our findings, SYN-AI was able to engineer genes from scratch by identifying evolution force associated with genomic building block formation as well as by applying natural selection protocols that mimic those that occur in nature. The evidence reported herein suggests that synthetic evolution methodologies are excellent tools for intelligent design of genes and should offer an alternative to rational design approaches. Notably, SYN-AI technology may be expanded allowing for design of genomes at very high resolution compared to current technologies that are based on the exchange of very large segments of genomic DNA. SYN-AI’s ability to write DNA code from scratch at high-resolution opens an endless potential for scientific exploration and gene design that may be applied to the evolution of any gene, dependent on the availability of PDB structural data. The ability to write DNA code at high-resolution also allows the rewiring of cell signal pathways. Saliently, SYN-AI synthetic evolution technology explores multiple evolution pathways based on the researcher’s experimental parameters, whereby each pathway results in formation of an alternate gene or gene family dependent on the mutability of the sequence space. As SYN-AI technology simulates evolution, outcomes also rely on randomness, whereby under identical experimental parameters there exists a possibility of exploring a diverse evolution pathway. Thusly, SYN-AI offers an excellent opportunity for the discovery of new chemistries that have potential applications in the treatment of cancer and other diseases as well as allow for the design of industrial genes.

## 4. Methods

### 4.1 High Performance Computing

SYN-AI was performed utilizing the Stampede 2 supercomputer located at the Texas Advanced Computing Center, University of Texas. Experiments were performed in the normal mode utilizing SKX compute nodes comprising 48 cores on two sockets with a processor base frequency of 2.10 GHz and a max turbo frequency of 3.70 GHz. Each SKX node comprises 192 GB RAM at 2.67 GHz with 32 KB L1 data cache per core, 1 MB L2 per core and 33 MB L3 per socket. Each socket can cache up to 57 MB with local storage of 144 /tmp partition on a 200 GB SSD.

### 4.2 Simulating Evolution

#### 4.2.1 Identification and Isolation of Genomic Building Blocks

SYN-AI analyzed evolution force associated with genomic building block formation across an orthologous sequence space comprising genes occurring at a homology threshold of > 80 percent identity to parental bovine brain 14-3-3 docking gene. The orthologous sequence space comprised 2.5 × 10^6^ bp of genetic material. Evolution force was analyzed by transforming the bovine brain 14-3-3 docking gene into a DSEC and performing 3 × 10^8^ DNA crossovers within genomic alphabets. DNA hybridization partners were randomly selected across orthologous sequence space. Evolution at the matter-energy interface was simulated by performing DNA hybridizations in a buffering solution of 3 mM Mg^2+^ and 1.2 mM dNTP at 328.15° kelvin. ^8^ Gibb’s free energy was calculated according to Owczarzy ^50^ and a penalty assessed for DNA base pair mismatches.

Evolutionarily fit DNA crossovers were selected by applying natural selection protocols. Neural networks limited selection to DNA crossovers based upon Gibb’s free energy. Genomic building blocks were passed thru pattern recognition filters that removed sequences displaying low sequence homology to the parental bovine brain 14-3-3 docking gene. Selection was limited to DNA crossover instances comprising evolutionarily favored mutations by quantum normalized Blosum 80 mutation frequency based neural networks. Natural selection was further accomplished by limiting selection to sequences characterized by (+) molecular wobble vectors. Subsequently, genomic building block libraries were constructed by quantum normalized neural networks that limited selection to DNA crossovers characterized by high magnitude of evolution force. Evolution force was enumerated over single and multidimensional evolution planes as described in ^71^.

#### 4.2.2 Evaluation of Evolution Force Associated with Genomic Building Block Formation

Evolution force associated with genomic building block formation was solved utilizing the rotation model as described in ^71^. Evolution force τ_*ϵ*_ was solved as a function of inertial moments *I*_*c*_ about the evolution axis and molecular wobble, **Eq. 1**. Whereby, inertial vector *I*_*c*_ is a function of evolution conservation *ϵ* and variance r from a recombinant pool comprising a total 3 × 10^8^ DNA crossovers, **Eq. 2**. Evolution conservation *ϵ* was solved as a function of DNA and protein evolution vectors 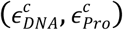, **Eq. 3**, and molecular wobble *ω*_*m*_ likewise solved as a function of evolution vectors 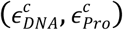, **Eq. 4**. The aforementioned evolution vectors are functions of DNA and protein similarity vectors (*X*_*i*_, *X*_*j*_) weighted by the recombinant pool, **Eqs**. (5 and 6). Whereby, evolution weights (*W*_*d*_, *W*_*p*_) describe similarity of the recombinant pool to the parental sequence in respect to DNA and protein primary sequence. Evolution weights are a function of mean DNA 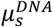and mean protein 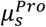 evolution vectors, **Eqs**. (7 and 8). Wherein,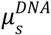 and 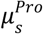 were solved by summation of genomic building block (GBB) similarity functions (*X*_*i*_/*n, X*_*j*_/*n*) occurring across the orthologue sequence space (*sspacece*^*r*^) divided by the total number of DNA crossovers *N*. Where, *sspacece*^*r*^comprised 2.5 × 10^6^ bp of genetic material.

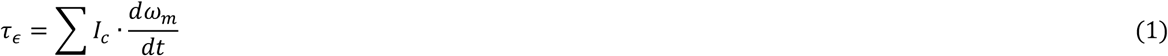

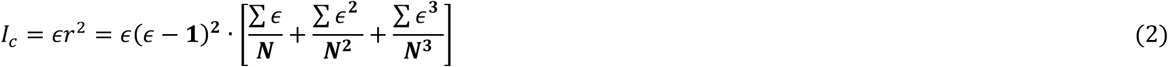

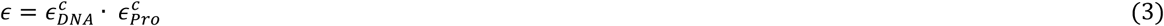

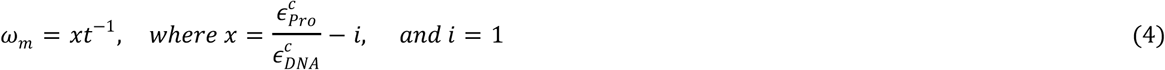

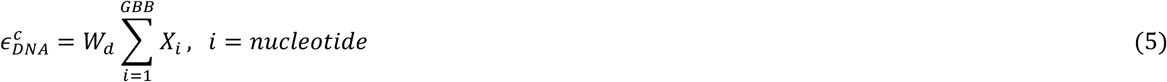

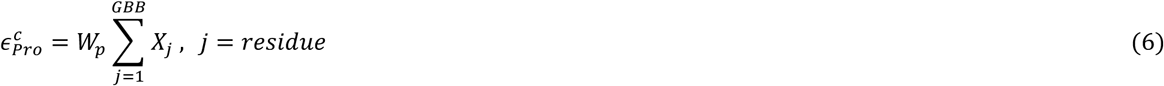

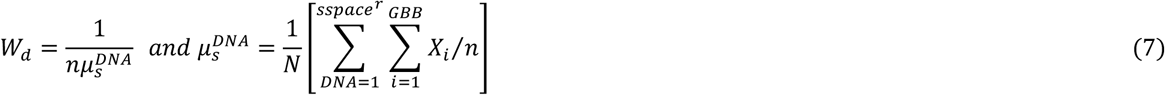

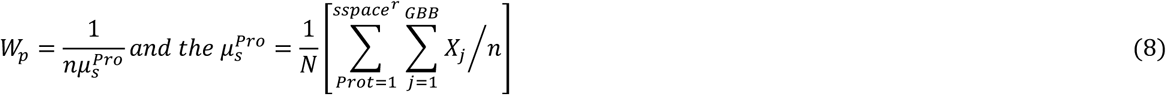

#### 4.2.3 Engineering Synthetic Super Secondary Structures

Parental super secondary structures were identified by partitioning the bovine brain 14-3-3 gene into a DNA tertiary code followed by analysis utilizing STRIDE ^51^ knowledge based secondary structure algorithms. Evolution was performed by ligation of genomic building blocks randomly selected from genomic alphabet libraries encompassing 5’ and 3’ terminals of parental structures. A cleaving algorithm was utilized to remove 5’ and 3’ prime overhangs from synthetic super secondary structures. Natural selection was performed by limiting selection to naturally occurring mutations utilizing Blosum 80 mutation frequency algorithms. SYN-AI also accomplished natural selection by imposing a secondary structure homology threshold > 90 percent identity, wherein synthetic sequences were aligned with parental 14-3-3 secondary sequences. A standalone version of PSIPRED 4.0 ^52^ was utilized to evaluate secondary structure. Synthetic super secondary structures were stored in DTER libraries for writing DNA code.

#### 4.2.4 Writing DNA Code from Scratch

DNA code was written from scratch by walking the DTER followed by random selection and ligation of synthetic super secondary structures stored in genomic alphabet libraries. SYN-AI constructed a library of 1 *X* 10^7^ genes that was passed thru a set of neural networks that evaluated closeness of synthetic protein structural states to native states. Wherein, SYN-AI set a minimal closeness threshold of > 90 percent identity according to ^71^. A subsequent selection limited super secondary structures to those characterized by naturally occurring mutations. The aforementioned was performed utilizing BLOSUM80 ^53, 54^ mutation frequency algorithms. A further round of natural selection restricted selection to synthetic 14-3-3 docking proteins characterized by mean secondary structure identities within the top quantile. Structurally conserved and functional 14-3-3 docking proteins were selected by a final natural selection protocol that evaluated closeness of protein active sites and hydrophobic interfaces to the parental bovine 14-3-3 docking protein. Wherein, selection was limited to synthetic proteins characterized by closeness thresholds of > 90 percent identity.

#### 4.2.5 Analysis of Synthetic Proteins

Sequence homology of synthetic proteins to parental bovine 14-3-3 docking protein was analyzed utilizing the Clustal Omega multiple sequence alignment tool.^12^ Wherein, phylogenetic analysis was performed utilizing Phylogeny.fr. Furthermore, we analyzed synthetic protein three-dimensional structure utilizing the I-TASSER suite.^55^ Wherein, protein-protein interaction and ligand-binding sites were analyzed utilizing Cofactor and Coach.^56^

## Acknowledgements

This work used the Extreme Science and Engineering Discovery Environment (XSEDE), which is supported by National Science Foundation grant number ACI-1548562. The authors acknowledge the Texas Advanced Computing Center (TACC) at the University of Texas at Austin for providing HPC resources that contributed to research results reported within this paper. URL: http://www.tacc.utexas.edu

---

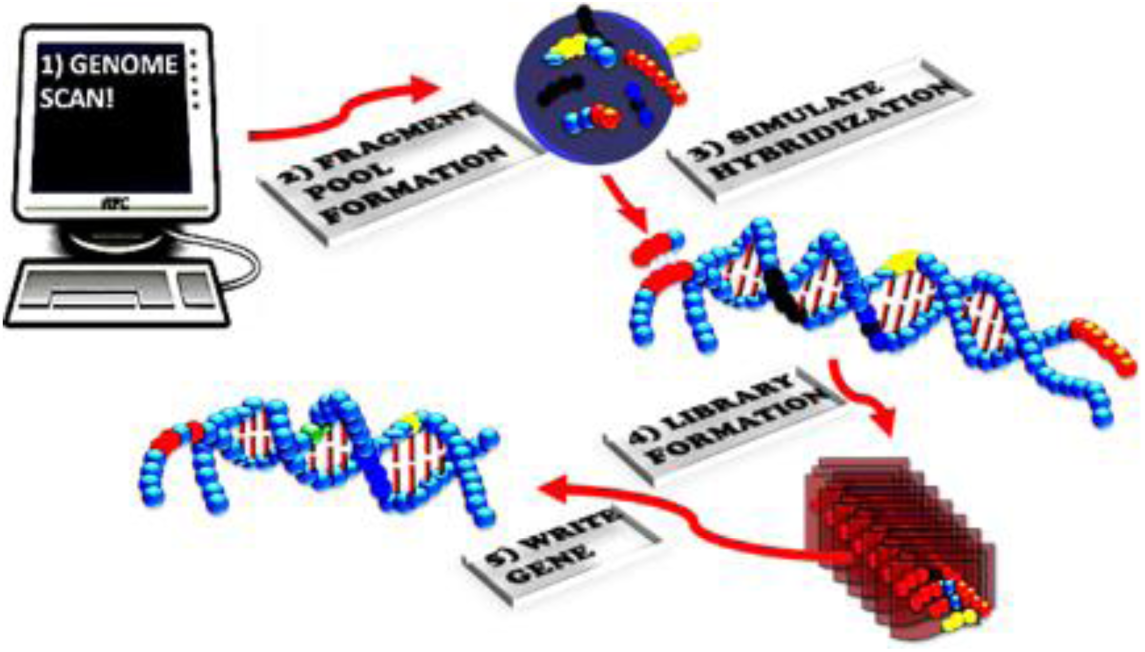

